# Monolayer graphene-on-polymer dressings promote healing and stabilize skin temperature on acute and chronic wound models

**DOI:** 10.1101/2021.05.16.444337

**Authors:** Marion Le Gall, Vincent Serantoni, Hervé Louche, Franck Jourdan, Dominique Sigaudo-Roussel, Christelle Bonod, Sandra Ferraro, Riadh Othmen, Antoine Bourrier, Latifa Dahri-Correia, Charlotte Hurot, Luc Téot, Vincent Bouchiat, Alain Lacampagne

## Abstract

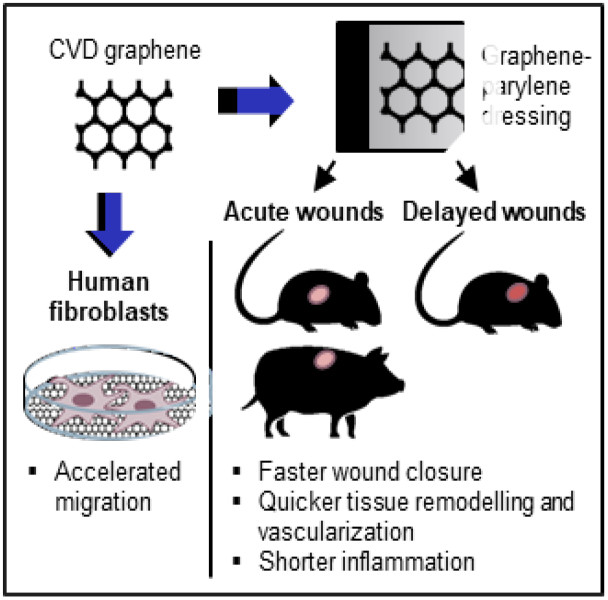

Monolayer graphene presented on the wound bed is assessed for its healing properties using both in vitro and in vivo models. For in vivo study, a cutaneous excisional wound is created on the dorsal surface of healthy and type-1 diabetic mice to mimic acute and delayed wound healing, respectively. A pig model is also chosen for its resemblance to human skin. Photographic and histological assessment of the wound are coupled with thermographic data recorded with an infrared camera. Graphene monolayer accelerates early phases of wound healing in vivo in every tested model. Upon removal of the bandage, wounds coated with graphene are less prone to temperature drop compared to the control samples. We hypothesize that graphene may directly shorten the inflammatory phase and/or enhance angiogenesis and cell migration in proliferative phase as demonstrated in vitro. Thermographic assessment of wounds could be of particular interest to follow both phenomena in an objective, rapid and non-invasive manner.

## 1. Introduction

Dressings are among the simplest and the most prevalent medical devices. Their common intent is to keep an open wound clean, free of external pathogens and dust and to promote its healing. While a lot of progress has been made in the polymer materials forming its main structure, there is still a need to improve the active material in direct contact with the wound bed, especially to control the volume of exudate, the presence and composition of a biofilm and to prevent the wound to turn into a chronic one.

Chronic wounds affect millions of people worldwide and lead to infections and amputations with devastating effects on life expectancy. The populations at risk are rising, including the diabetics and the elderly. In 2015, between 2.4 and 4.5 million of Americans suffered from them.^[1]^ Chronic wounds have become a global health problem, putting a huge burden on healthcare economy. Clinically, they are defined as wounds older than 4 weeks, usually suspended in the inflammatory phase of healing.^[2]^ The factors leading to chronicity are multiple and complex. Notably, too many neutrophils at the wound site cause an excess of inflammatory markers, which degrade the growth factors necessary to enhance the proliferation and differentiation of fibroblasts and keratinocytes. The vascularization is also impaired due to the deregulation or resistance to different growth factors.^[3–5]^ Infection is also known as an important player in the shift between acute and chronic wound.^[6,7]^

Many innovative dressing materials have been considered to improve patients’ prognosis and quality of life, among which the family of carbon-based materials at the crossroads between organic and inorganic materials seems promising.^[8]^ In particular, Graphene is a two-dimensional atomically-thin crystal of pure carbon atoms, covalently bound into a honeycomb lattice. Isolated 15 years ago and recently mass-produced, it features an outstanding combination of physical and chemical properties with promising applications in many technological fields. In the biomedical domain, graphene’s biocompatibility and high electrical conductivity make it an interesting candidate for biosensing or tissue engineering.^[9–13]^ Furthermore, the anti-coagulant, anti-bacterial and electrophysiological effects of graphene could help to meet some of the persistent challenges in wound healing,^[14–17]^ for example regarding chronic wounds. Recently, nanofibers of graphene were directly applied to excisional wound on mice and rabbit model to assess their impact on the wound healing rate.^[12]^ Closed wounds were visually achieved quicker with a graphene/chitosan material than without graphene and without dressing, most probably thanks to an anti-microbial effect. However, the acute model used in this study was not relevant to human as the skin healing in rodents uses mainly contraction of the skin and subcutaneous tissues whereas human skin needs to create a new granulation tissue to replace the damaged area.^[18,19]^

The present study aims at investigating the effect of a graphene monolayer-based dressing on wound healing parameters. Several models were used to best approximate the healing behavior of human skin: human dermal fibroblasts *in vitro*, healthy or type-1 diabetes mice to model acute or delayed wound, respectively, and healthy pigs. Different indicators were chosen for wound healing, including an original non-invasive thermographic methodology.

## 2. Experimental section

### Graphene synthesis and transfer

Graphene layers were obtained by low-pressure chemical vapor deposition (CVD). Graphene was produced by catalytic decomposition of methane gas CH_4_ at 1,000°C in diluted hydrogen atmosphere on 25-μm-thick copper foils (99.8% purity, Alfa-Aesar). After surface cleaning in acetone, the copper foils were placed in the reactor oven, annealed in diluted H_2_ atmosphere (dilution in Ar at 10%) at 1000°C for 2 hours. This thermal treatment helped to reduce the native copper oxide and enlarge the copper grains. Then, pulses of methane gas CH_4_ (2 sccm 10 s, then 60s off) were injected in the oven chamber as a carbon source, instead of using continuous flow of methane, to prevent the aggregation of carbon at the nucleation centers and avoid the formation of multi-layers patches. This resulted in a continuous, polycrystalline graphene layer conformably covering the metal layer.

For the studies on glass substrates, a classical poly(methyl methacrylate)-based graphene transfer method was used.^[20]^ For the studies on biocompatible parylene C, the graphene ultrathin layer on copper was covered by a 10 *μ*m-thick polymer film by gas deposition. Then, etching of the catalytic graphene copper film was performed in aqueous Sodium Persulfate solution (100g/l). The resulting Graphene-on-polymer film was then thoroughly cleaned in deionized water to remove all traces of copper residues.

### Cell Culture

Human dermal fibroblasts from a breast skin biopsy of a 20-years-old woman were grown in DMEM-F12 Ham medium (Sigma-Aldrich) supplemented with 10% fetal calf serum and 1% ZellShield™ (Minerva Biolabs, Germany), at 37°C in a 5% CO_2_ incubator.

### Scanning electron microscopy (SEM)

Glass slides of 12 mm diameter were inserted in a 24-wells sterile plate. Some were coated with a CVD graphene monolayer, whereas the plain glass surface of others was used for control samples. Fibroblasts were counted and 28 000 were seeded in each well. Cells were grown for 72 hours, fixed in a solution of glutaraldehyde 2%, cacodylate 0.18 M, stabilized at pH 7.2 and then dehydrated in ethanol (from 30°C to 100°C) and 100% HMDS (Sigma). The slides were metallized by gold physical vapor deposition in high vacuum. Other images of cells on graphene were made without metallization thanks to this material’s electrical conductivity (**Figure S1.**). High resolution SEM imaging was performed using a 1keV Zeiss Ultra Field Emission Gun Scanning Electron Microscope.

### Immunofluorescence

Immunofluorescence was performed on glass slides of 12 mm diameter, with or without graphene coating, previously inserted in a 24-wells sterile plate. 3800 human dermal fibroblasts were counted and seeded in each well. Fibroblasts were grown during 48 hours, fixed and permeabilized 5 minutes at −20°C in presence of 100% methanol. After the blocking phase in 1% BSA, cells were incubated overnight, at 4°C, in presence of primary antibodies against vimentin (Sigma-Aldrich V6630). After three washes in PBS 1X, they were incubated 1 hour with the fluorescently-conjugated secondary antibodies anti-mouse Alexa fluor 488nm (Thermo Fisher A-11029). The staining of F-actin was performed by fixing the cells during 15 minutes in 4% PFA at room temperature, permeabilizing for 10 minutes in 0.1% triton X-100 and incubating with phalloidin-TRITC (Sigma-Aldrich) for 1 hour at room temperature. Nuclei were stained for 5 min in DAPI solution. Coverslips were mounted with Permafluor (Thermo Fisher, TA-030-FM). Images were captured using an Eclipse Ti-E 300 inverted microscope (Nikon) with a collsnap fx CCD camera (Photometrics, USA) and Metavue software (Universal Imaging Coporation, USA). Image analysis was performed using imageJ Software (National Institut of Health, USA).

### Proliferation assay

Proliferation tests were performed using 24 wells sterile Plate. 3800 cells were counted and seeded in each well on a previously inserted 12 mm diameter glass slide coated with or without graphene. Each condition was repeated three times. Cells were then counted with trypan blue each day for 9 days.

### In vitro graphene 2D migration assay

An *in vitro* migration assay was elaborated by designing gaps in between cell cultures. A direct mechanical scratch of a cell monolayer was not possible as it could have damaged the fragile graphene layer. Therefore, elastomer molds (Ibidi® cells) were used. They consist of two adjacent reservoirs separated by a 500 *μ*m-thick elastomer wall in contact with the plate. Each well of a 24 wells sterile plate was filled with a single Ibidi® cell and about 15 000 cells were seeded in each reservoir. Fibroblasts were cultured for 48 hours until they adhered and formed a monolayer. The elastomer molds were then removed, leaving two defined cell patches separated by a 500 *μ*m-wide gap. Cells were rinsed three times with PBS. Then, PBS was replaced by DMEM-F12 Ham medium containing a reduced percentage of fetal calf serum (1% versus 10% in the culture medium) to limit proliferation. Cell migration was monitored along the gap and captured with an inverted microscope 16, 24, 38 and 48 h after seeding. Migration quantification was processed using Image J and the plugin “cell tracking”. Three independent experiments were carried out.

### Mouse model

All experiments were conducted in accordance with a protocol approved by the Institutional Animal Care and Use Committee of Languedoc–Roussillon (n°2018013109035735). Two mouse models were used in this project.

(1) Delayed wound model (n=20): 5 weeks-old, fasting males C57BL/6j mice (Janvier Labs, France), were injected with an intraperitoneal bolus of streptozotocin (S0130, Abcam) at 180 mg/kg in citrate buffer (pH 4.5). 48 hours later, the mouse glycaemia increased to 300 mg/dL if type 1 diabetes was induced. Then insulin was injected intramuscularly twice every week. 4 weeks later, mouse back was gently shaved.

(2) Acute wound model (n=20): the lower back of 9 weeks-old male C57BL/6j mice (Janvier Labs, France), were shaved.

The day following shaving, the mice from the two models were anesthetized under 3% Isoflurane and a skin biopsy was sampled using an 8mm diameter biopsy punch. Anti-contraction splints were set up. One type of dressing (graphene-coated parylene or graphene-free parylene (SHAM)) was randomly assigned prior to the beginning of the experiment and applied to each mouse.

Dressings were applied directly in contact with the wound bed to cover it whole surface. Surgical KwikSill© glue kept the dressing and the anti-contraction splint in place. For the acute wound, cutaneous wound temperature (CWT) values were recorded on day 2, 4, 6 and 8 before removing the dressing, after dressing removal and after applying a new dressing. Planimetry photographs were taken at each stage. On day 10, only temperature and photographic measurements were performed, no new dressing nor splint were applied. On day 16, blood and skin biopsy over the wound area were sampled. For histological analysis of wound healing phases, 2 mice per group (SHAM or graphene) were sacrificed at day 4 and day 10. For the chronic wound, CWT values were recorded at day 2, 5 and 9, before removing the dressing, after dressing removal and after applying a new dressing. At day 12, only temperature and photographic measurements were performed, no new dressing nor splint was applied. At day 21, blood and skin biopsy over the wound area were sampled.

### Pig model

All experiments were conducted in accordance with a protocol approved by the VetAgro Sup Ethical Committee (n°1511_v2). Eight skin excision of 4×4 cm were created on the back of three male Youna pigs, 50 kg (GEAC des 4 vents, France), anesthetized with Tiletamine and Zolazepam (Zoletil 100®) 5mg/kg. Each wound was randomly covered with a parylene dressing coated with graphene, or not (SHAM). The dressing was applied directly on the wound bed and kept in place with damp pads, a semi-occlusive bandage (Tegaderm®), a dry compress and Tensoplast® around the animal body. At day 3, 7, 10, 14, 17, 21, 24, 28, 31 and 35 the dressings were removed, the wound was washed with saline serum solution and a new dressing of the same group was re-applied. Photographic assessment was carried out at the same time. For histological analysis of wound healing phases, one pig was sacrificed at D10, one at D17 and one at D35.

### Photographic assessment

Photographic assessment of the excisional wound was performed using magnifier Leica M60 and IC80D. Photographic planimetry of the wound was repeated three times, using ImageJ (ver.1.48v, Wayne Rasband). The wound percentage of closure was calculated as the ratio between the wound area of the calculation day and the wound area at day zero.

### Histological assessment

Mouse skin biopsies were fixed in PFA 4% overnight then washed with PBS and incubated in glucose baths before inclusion in Tissue Tek© O.C.T., and congelation in isopentane cooled down with liquid nitrogen.^[33]^ Pig skin biopsies were fixed in AFA (Alcohol-Formol-Acetic acid liquid) for 24h at room temperature, dehydrated and paraffin-embedded. 7*μ*m (mouse) and 5*μ*m (pig) transverse sections of skin and subcutaneous tissue were cut using a cryostat. Tissue sections were stained with hematoxylin, eosin and saffron. Slides were examined with an optical microscope (Leica DM2000) and a camera (Leica DFC420C), using the acquisition data software LAS (ver. 4.2). Epidermis thickness was measured in segmented pictures. An average of at least 30 measurements was computed using ImageJ (ver.1.52g, Wayne Rasband), as the epidermis thickness per sample.

### Cutaneous wound temperature assessment

CWT was measured with a thermic infrared camera Cedip W (Titanium 640×512 px) calibrated with a black body. The temperature of the wound area and peri-lesional skin (3284 pixels) was measured every second during 4 minutes. During the first 2 minutes, the wound was covered by the dressing then, the dressing was removed and the temperature varied for another 2 minutes.

The mean temperature variation was normalized with the maximum temperature accounting for 100%. The standard deviation between the pixels after dressing removal was also calculated. The thermal imaging recordings were analyzed by using Altair (ver. 5.91.010, FLIR Systems™), Matlab® (ver.2014b, MathWorks) and Prism (ver. 7.0, GraphPad).

### Blood analysis for cytotoxicity assay

Mouse blood was sampled on day 16 for acute models and day 21 for delayed models. Serum from was retrieved after 10 min, 2000 g centrifugation then stored at −80 °C. C-reactive protein (CRP, Mouse CRP assay ref MCRP00, R&D Systems®) and creatine kinase (CK, creatinine assay ref 65340, Abcam®) were measured with ELISA kits as per manufacturer’s instruction.

### Statistical analysis

Statistical analyses were performed with Prism (ver. 7.0, GraphPad). Data are presented as mean ± standard error of mean. When appropriate, one-way ANOVA was used to compare groups with correction for multiple comparison. To analyze the temperature mean variation, a two-way ANOVA was used with a Bonferroni correction for multiple comparisons. Significance was fixed at a *p* value < 0.05. All analyses were blinded.

## 3. Results

The graphene used in this study was obtained by CVD of methane gas on a copper foil. This technique inherited from the microelectronics industry results in a continuous, polycrystalline graphene layer growth on the metal.^[21]^ To establish its interest in wound healing applications, the graphene monolayer was subsequently transferred on a non-toxic, electrically insulating substrate (glass or polymer), and the copper foil removed by chemical etching.

### 3.1. *In vitro* study

In a first step, human dermal fibroblasts were grown in well plates decorated with glass slides coated with graphene or not (control). SEM images in **Figure S1.** proved the integrity of the graphene coating after cell seeding. Interestingly, folds on the graphene surface were observed close to the cells’ filipodia. A series of assays was then led for a better understanding of the effect of the graphene monolayer on the cells *in vitro*.

First, proliferation assays gave similar results on graphene or on bare glass surface, reaching an equal number of cells on day 9 (D9) (**Figure 1. a.**). There was a slight peak decrease at D8 with graphene (p=0.001 ***).

**Figure 1.**
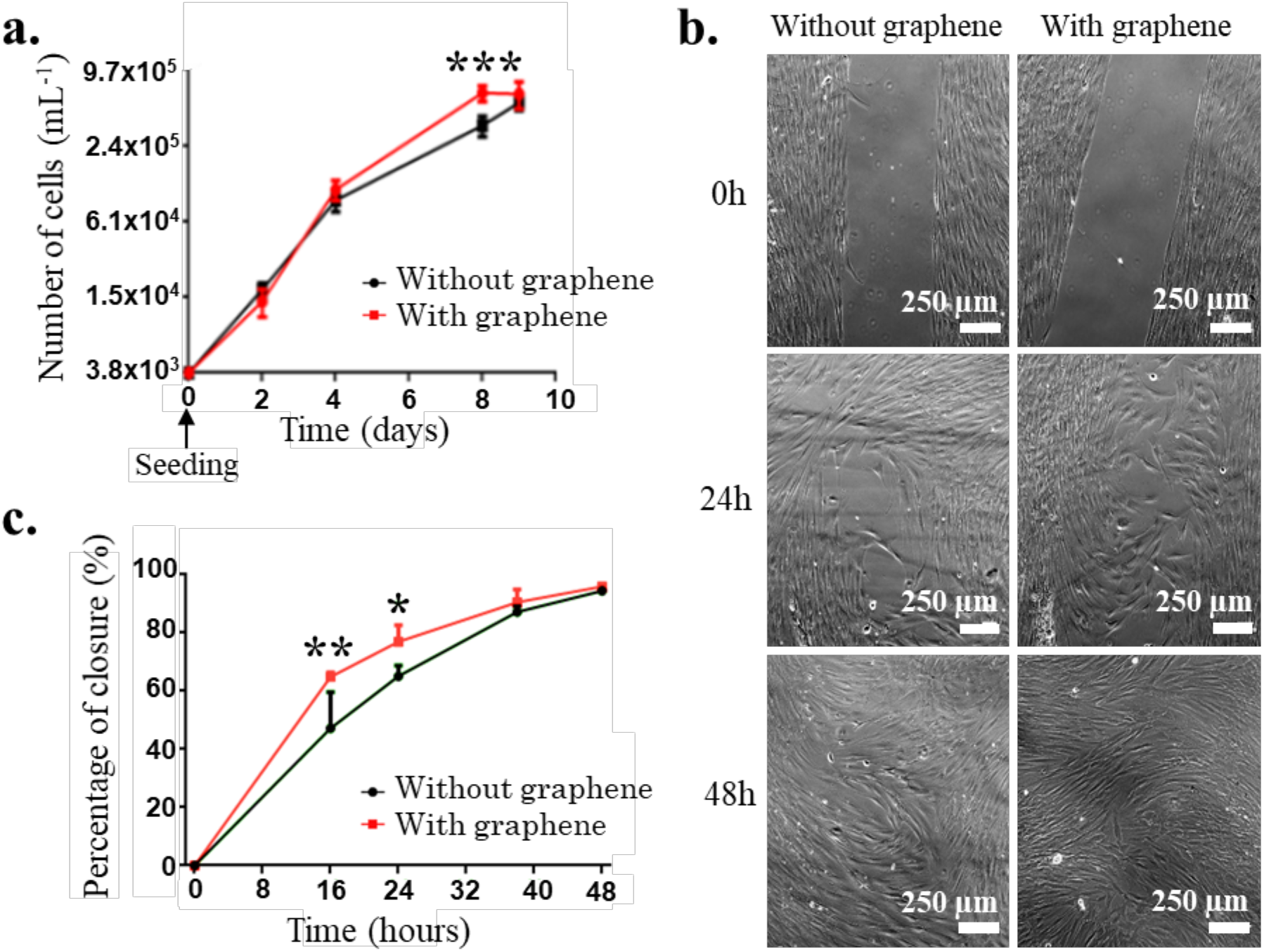
2D migration of cultured fibroblasts is increased on graphene. **a.** Growth curve analysis of dermal fibroblasts from woman breast skin biopsy in the presence (red) or absence (black) of graphene. **b.** Images of migration tests on fibroblasts seeded on graphene (right) or not (left), recorded after 0h, 24h and 48h. **c.** Calculated percentage of gap closure during migration assay. All data are shown as mean ± standard error, * p < 0.05, ** p < 0.01, *** p < 0.001 vs SHAM.

The SEM images showed that the morphology of the fibroblasts was similar on both substrates (**Figure S2. a.**). This observation was confirmed by immunofluorescence (**Figure S2. b.**). The cytoskeleton of the cells was not impacted by graphene, actin and vimentin remained well organized with well-formed stress fibers and intermediate filaments.

Furthermore, cell migration was evaluated in the presence and in the absence of graphene (**Figure 1. b.**). In the initial stage, after insert mold removal (see methods), the borders between the acellular zone and the confluent monolayers were clear and the cells parallel to the border. After 24h, the migration of the fibroblasts seemed to be more homogeneous on graphene. To get a quantitative assessment of this observation, we calculated the ratio between the surface colonized by the cells at the calculation time and the gap surface at the initial time (**Figure 1. c.**). This unraveled a significantly higher percentage of closure after 16 hours (p < 0.01) and 24 hours (p < 0.05) with graphene compared to bare glass. Cell confluence was reached after 48 hours in both groups.

### 3.2. Preliminary considerations towards *in vivo* studies

Thereafter, parylene C was laid on top of the single graphene layer and the copper foil was chemically destroyed to recover the dressing made of graphene on a parylene substrate. Parylene C is known for its biocompatibility and widely used to coat implantable devices. In the following parts of this study, graphene-coated dressings and parylene-only dressings (SHAM) were deposited directly on wound beds in different models, to cover the whole surface of the wound.

It is important to note that the graphene used in this study is a pure crystalline monolayer of carbon as opposed to graphene oxide in solution. Graphene oxide is a derivate of graphene that has been reported as possibly toxic to bacteria but also triggering cellular damage in its solution suspended form. ^[14,22,23]^ However, in the absence of oxidation, graphene should be biologically passive. As a precocious measure, the blood concentration of CRP or CK, two main proteins involved in a general inflammation response and cell lysis, were measured and compared in SHAM and graphene-dressing populations after 10 days of care (**Figure S3**). No significant difference was observed, confirming the absence of cytotoxicity of the studied material.

The improved wound healing provided by graphene has been demonstrated and suggested to be caused by its anti-bacterial effect.^[12,24,25]^ Consequently, bacteria presence was tracked in our laboratory on the intact skin and wound area of C57BL6/J mice. Some *streptococcus* (*sciuri* and *xylosus*) in low quantity, and some *bacillus* and *enterococcus faecalis* on fewer samples were detected after seeding exudate in supplemented medium up to 3 days (data not shown). Therefore, bacteria population was very low on our laboratory mice and any observed beneficial effect of graphene was unlikely to be related to its anti-bacterial properties.

### 3.3. Acute wound model

An excision wound model of healthy C57BL6/J mouse accounted for acute wound.^[26]^ Each mouse was assigned a dressing randomly (graphene or SHAM) that was changed every 2 days for 10 days. The dressing composition is depicted in **Figure S4. a.**. Anti-contraction splints have been used to overcome the contractions of the skin induced during during the healing of wounds in the mouse models and thus to come closer to the processes of human healing.

The efficiency of graphene and SHAM dressings to accelerate wound healing were compared based on several indicators. Classically, photographic and histological assessments were used to observe tissue evolution and to calculate the percentage of wound closure and the epithelial thickness.

The photographic follow-up of injuries (**Figure 2. a.**) revealed no significant macroscopic change of wound healing speed during the first 16 days post wound creation. Quantitatively, the measured wound closure did not differ between the groups (at D2 *p*=0.55, n=10).

**Figure 2.**
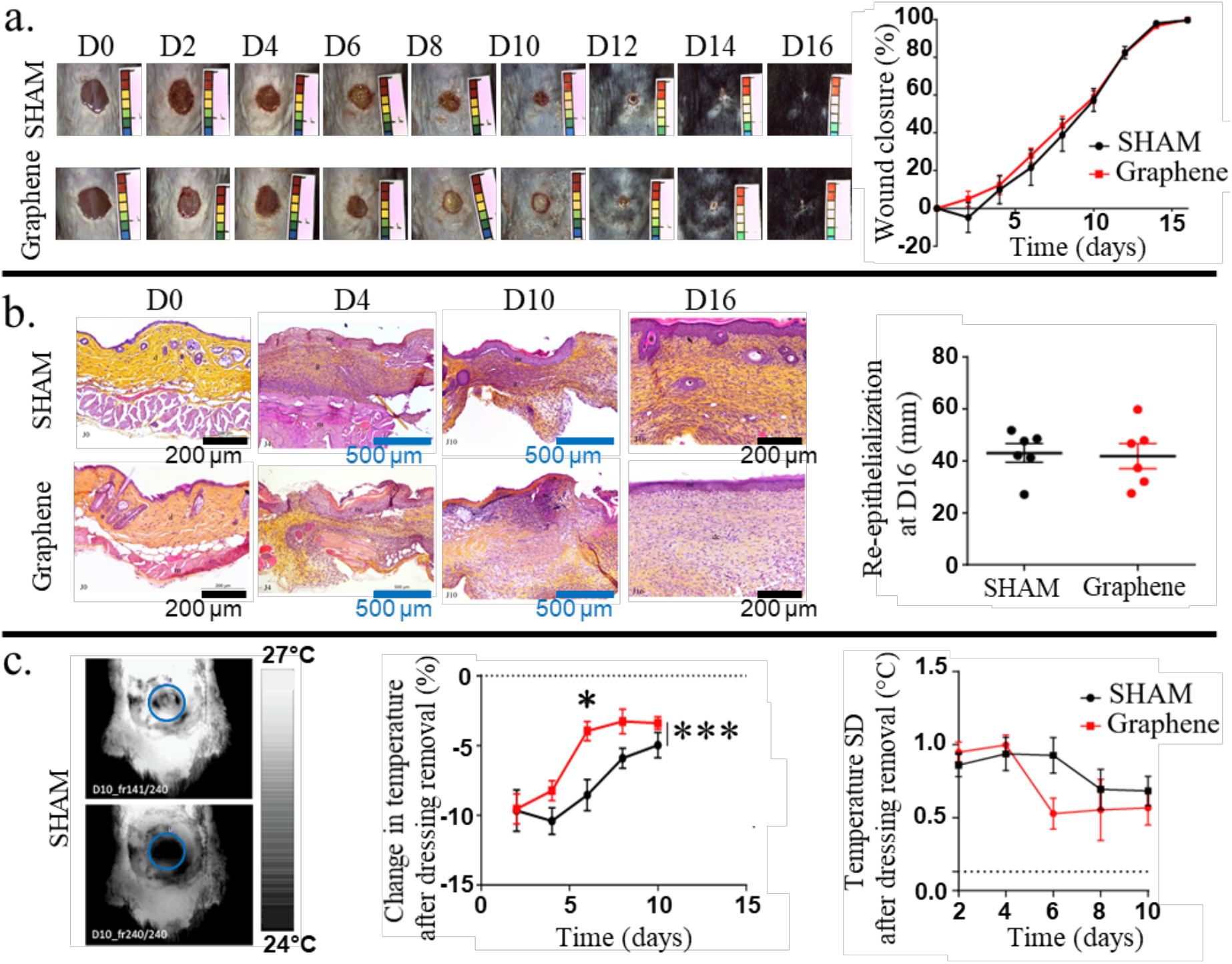
Early boost of healing on a mouse acute model wearing graphene dressing. **a.** Representative photographs of the wound evolution for SHAM and graphene groups (left). Quantification of wound percentage of closure through time (right). **b.** Representative hematoxylin eosin saffron-dyed images of wound bed histology in SHAM or graphene groups (left). Dot plot of the re-epithelialization (epidermis thickness) at D16, n=6/gp (right). At D4 (n=2/gp), the neo-epithelialization of the wound dressed with graphene was more advanced. There was less granulating tissue and more differentiated cells compared to SHAM. The phase at D10 (n=2/gp) and D16 (n=6/gp) does not differ between the 2 groups. **c.** Representative thermographic picture after dressing removal (top left) and 100 s later (bottom left) at day 10 on a SHAM mouse. The blue circle locates the wound area. Quantification of the change of normalized surface skin temperature of the mouse after dressing removal (center). Quantification of the temperature standard deviation (SD) at the wound area after dressing removal. The horizontal dot line is the standard deviation of the skin temperature before creating the wound (right). All data are shown as mean ± standard error, * p < 0.05, |*** p < 0.001 vs SHAM on the global curve.

However, the histological assessment gave attention to some variations as early as D4 (**Figure 2. b.**). In SHAM group, necrotic and inflammatory tissues were observed close to the skin, with an inflammatory and vascularized matrix near the muscle. In contrast, with graphene dressing, a thick and non-organized neo-epidermis characterized the wound, with a subjacent underdeveloped extracellular matrix and the presence of macrophages and polynuclear cells. As the wound healing progressed, at D10, the re-epithelialization process started in SHAM biopsies and a granulation tissue developed underneath. The new blood vessels were dilated. Meanwhile, the re-epithelialization in graphene biopsies was nearly complete. The subjacent extracellular matrix was hyper-vascularized with persistent inflammatory cells. After healing on D16, there was no histological difference between the two groups. The re-epithelialization was complete with a differentiated, organized and layered neo-epidermis. Quantitatively, the final epidermis thickness was the same with both dressings.

In the present study, we propose another indicator of wound healing based on temperature measurements. Indeed, temperature is known to be an important factor during healing^[27–29]^ and thermography is already a reliable and valid tool to evaluate burns.^[30,31]^ Cutaneous temperature change is correlated to arterial stenosis^[32]^ and detects acute limb ischemia.^[33]^ Thermography measurement records the hyper-vascularity resulting from tumor.^[34]^ Here, the mean temperature of the wound area and peri-lesional skin was measured (1 frame/s, 3284 pixels). During the first 2 minutes of recording, the wound was covered by the dressing, then the dressing was removed and, as expected, the temperature at the wound area decreased. Two quantitative indicators were calculated from this data: the percentage of change in the mean temperature and the standard variation of the temperature between the pixels in the recording area and during the first 120 s after dressing removal.

We addressed how the cutaneous wound temperature changed throughout healing (**Figure 2. c.**). There was a significant variation between the two populations (*p* < 0.001, n ≥ 8).

In SHAM group, there was a moderately higher decrease in CWT when the dressing was removed on D2 than after wound creation. Afterwards as the wound healed, this variation linearly dropped. In comparison, with graphene dressing, the variation of temperature after dressing removal was systematically lower, with a significant difference on D6 (*p*=0.011, n=8).

Regarding temperature homogeneity, a 2-fold decrease of the standard variation was measured between D4 and D6 for wound covered by graphene.

### 3.4. Delayed model

The purpose of a delayed model in this study was to assess if graphene improves healing in a pathological situation where oxidative stress and vascularization are impaired. A well-described model was used, by inducing type 1 diabetes in C57BL6/J mice with a single streptozotocine injection.^[35–37]^ Excision wounds were dressed with or without graphene for 12 days. In this model, healing was indeed slower, and exudates were more voluminous. The dressing change was adapted accordingly (D2, D5, D9 and D12).

Photographic wound tracking suggested a visual reduction of the wound in presence of graphene starting from D5 until dressing final removal at D12 (**Figure 3. a.**). Indeed, the calculated wound closure was greater at D5 (*p*=0.067, n=10) and D9 (*p* > 0.05, n=10) with graphene dressing compared to SHAM. On D12, the shift between the groups decrease to end at the same closure rate at D21.

**Figure 3.**
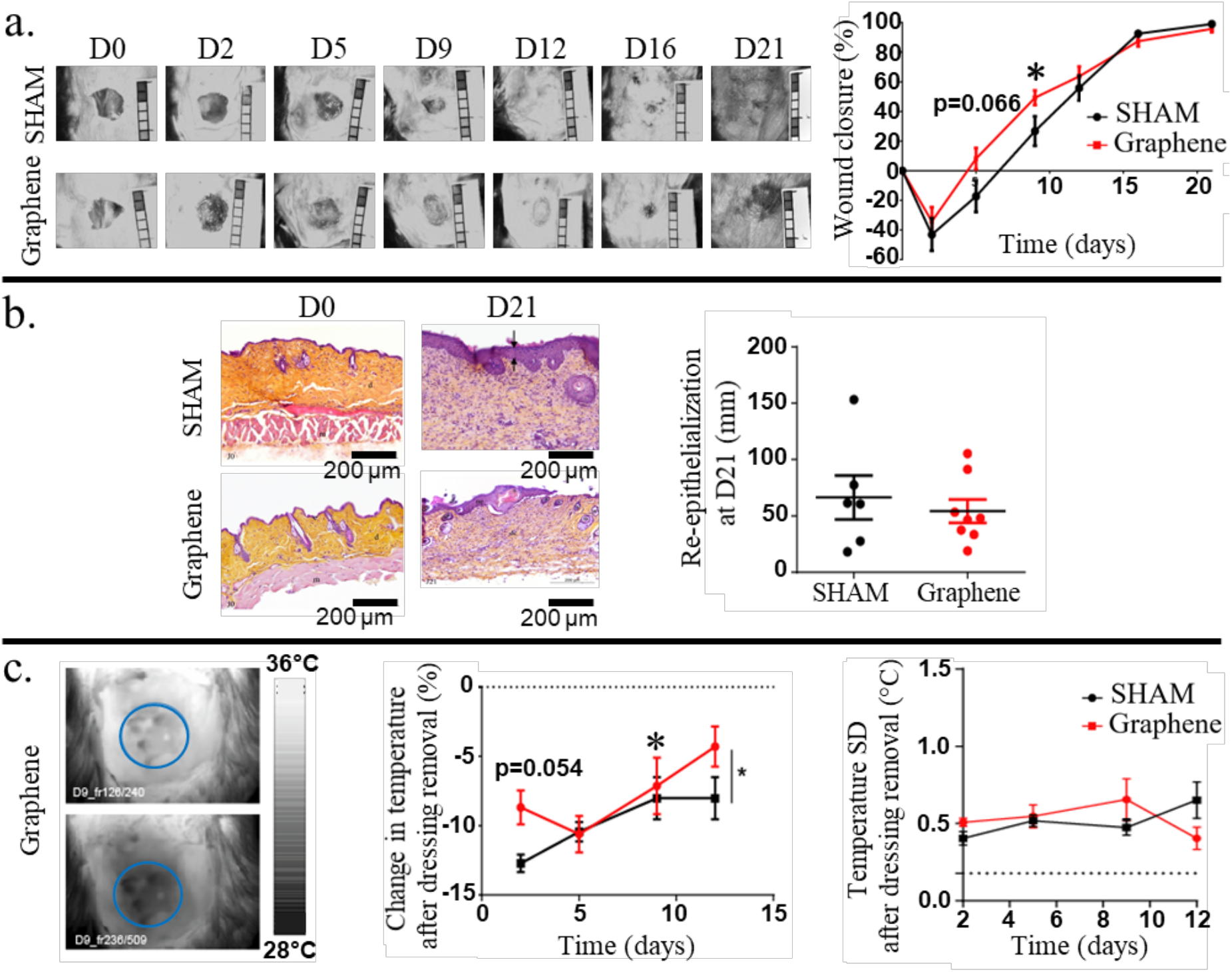
Graphene improves healing early phase on a mouse delayed wound model. **a.** Representative photography of the wound evolution for SHAM and graphene (left). Quantification of wound percentage of closure through time (right). **b.** Representative hematoxylin eosin saffron-dyed histology image at day 0 and 21 after wound excision (left). Dot plot of the re-epithelialization (epidermis thickness) at D21, n ≥ 6/gp (right). **c.** Representative thermographic picture of the dorsal area of the mouse just after the dressing removal (top left) and 110 s later (bottom left) at day 9 on a graphene mouse. The blue circle locates the wound area. Quantification of the change in normalized surface skin temperature after dressing removal (center). Quantification of the temperature standard deviation (SD) at the wound area after dressing removal. The horizontal dot line is the standard deviation of the skin temperature before creating the wound (right). All data are shown as mean ± standard error, n ≥ 7 animals/group. *p < 0.05 vs SHAM, |*p < 0.05 vs SHAM on global curve.

Skin biopsies histological assessment on D21 did not reveal any difference between the two groups at the end of the healing process. The samples were in early remodeling phase with granulation tissue characteristics: hyper-cellularity, major neo-vascularization and constant inflammatory response (**Figure 3. b.**). There was no difference in skin thickness.

Regarding CWT in **Figure 3. c.**, there was a significant difference depending on the dressing used (*p*=0.03, n≥6). In SHAM mice, CWT variation decreased linearly throughout the first 9 days of healing and reached a plateau at D9. In comparison, with graphene, the CWT changed less at all time points (at D2 *p*=0.54, n ≥ 8 and at D9 *p* < 0.05, n ≥ 6). The thermal distribution at the wound area after dressing removal was not different between the 2 groups and did not vary during wound healing.

### 3.5. Pig model

Pig skin has many more similarities to human skin than mouse skin and consequently provides a model closer to the target of the dressing we are using: people with diabetes. In consequence, we have investigated graphene-dressing effect on a cohort of 3 pigs in collaboration with a contract research organization (BioVivo, Bron, France). The dressing composition is depicted in **Figure S4. b.**.

According to photographic assessment, graphene dressing seemed to promote wound closure (*p*=0.0537 on D10, n≥3, **Figure 4. a.**). Due to the low number of animals processed, no statistical test was performed on the re-epithelialization measurement. However, the wounds covered with graphene dressings seemed to reach largest skin epidermis values compared to the wounds without graphene (**Figure 4. b.**).

**Figure 4.**
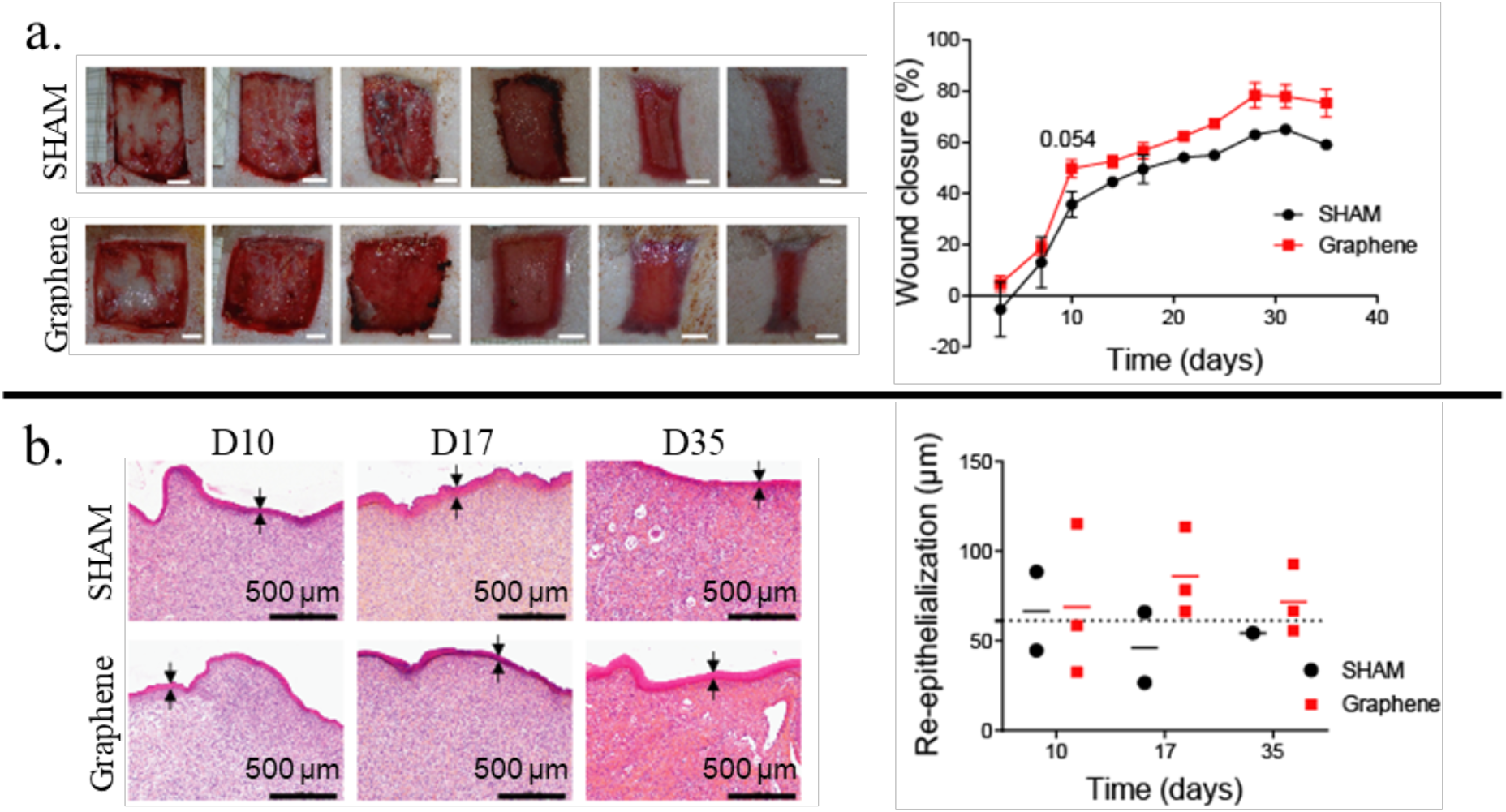
Graphene dressing promotes wound healing on an acute pig model. **a.** Representative photographs of the wound evolution for SHAM and graphene groups, scale white bar: 10 mm (left). Quantification of wound percentage closure through time (right). **b.** Representative hematoxylin eosin saffron-dyed histology assessment at day 10, 17 and 35 after wound excision (left). Dot plot of the re-epithelialization (epidermis thickness) at D10, D17 and D35. Dot line is the skin epidermis at D0 (right). All data are shown as mean ± standard error, n≥2 dressings/animal. * p < 0.05 vs SHAM.

## 4. Discussion

We investigated the potential of graphene dressings as wound healing enhancers. The graphene was produced according to an original pulsed CVD method on a copper foil.^[38]^ The graphene monolayer was transferred on a film of parylene C to fabricate dressings. Dressings with or without graphene were cut specifically to cover the whole cutaneous lesions with a direct physical contact between graphene layer and wound bed.

The *in vitro* study proved that the graphene monolayer was not damaged in the presence of cultured human dermal fibroblasts. This first result is crucial in view of transposing the material *in vivo* into dressings without the risk of releasing residues into the wound and while maintaining the effectiveness of the coating. In return, neither the proliferation nor the morphology of the cells was impacted by the presence of graphene *in vitro*. The relationship between cell migration capacity and cytoskeleton organization is well established. Here, not only was cell mobility preserved on graphene, but cell migration was even faster on this substrate than on glass, which was very promising for accelerated wound healing. This may be due to the softness of graphene, which can deform and provide a favorable 2D-matrix. This hypothesis is sustained by the observation of folds on the graphene surface, which could indicate that the cell is probing the surface stiffness thanks to its filipodia.^[39]^ This observation supports the hypothesis of Lasocka *et al*., who suggested a putative action mechanism of graphene on wound healing.^[40]^

Cohesively, the graphene dressing did not induce a systemic inflammation in healthy mice *in vivo*. In our acute model, we evaluated how graphene affects the wound healing kinetic. No difference was found regarding the percentage of wound closure between graphene and SHAM. However, histological assessment showed advanced wound healing phase in graphene group that tended to show that the wound in contact with graphene had a shorter inflammation phase. The neo-epidermis present in graphene on D4 wound bed indicated the beginning of proliferation stage while SHAM skin biopsy was still in the inflammatory phase with necrotic and inflammatory tissue. At D10, remodeling was properly started in the graphene group with a nearly complete re-epithelialization. These observations were coincident with thermal variations: there was a lower CWT decrease after dressing removal with graphene from D4 to D8, accompanied by a more homogeneous temperature distribution. This could reveal an earlier angiogenesis, as suggested in the aforementioned literature, or a downregulation of the inflammation response. Indeed, in cancer detection, cutaneous temperature change was associated to an inflammatory response and not to the hyper-vascularity of the tumor induced angiogenesis.^[19]^ Therefore, thermographic assessment could turn out to be a particularly relevant indicator of inflammation in wound healing. It is nondestructive and seems highly correlated to histology. Moreover, CWT monitoring gives complementary and objective understanding of the wound healing progress than wound photographs. Macrophages staining on histological slides or immunohistochemistry could be used to quantify the inflammatory response and offer a clear parallel with the decrease of temperature variation. Laser-Doppler assessment of microcirculation could be used in parallel.

The chronic wound was simulated by diabetes mellitus type 1, induced by streptozotocine. The faster wound closure (147% area difference on D5) in the graphene group suggested that graphene dressings accelerate wound healing in delayed model. Here again, the CWT measurements on the same time points most likely reflects a reduction of the inflammatory phase or a faster revascularization. However, as diabetes per se induced impairment of the redox and the vascularization system, CWT measurement appeared to be less discriminant in this model, especially in the early phase of healing.

To get closer to the behavior of human skin, pig models were also tested. Here again, the wound closure rate was higher with graphene dressings.

The anti-bacterial properties of the graphene are hardly tested while using experimental models in laboratory animal facility. Our animals have a very limited cutaneous microbiota due to their special handling and environment. Therefore, increased wound healing through anti-bacterial performance seems unlikely in our models. Based on the quickest migration of fibroblasts in the presence of graphene, which was not due to an increased proliferation, we hypothesized that the graphene may act as an electrically conductive network that would connect the cells and stimulate their migration. This would in turn reduce the inflammation phase duration, as it has been previously demonstrated with neuronal cells.^[41–43]^ We may also hypothesize that this improved cellular migration results in faster angiogenesis in the presence of graphene, as suggested by the histological analysis and change in CWT.

Further tests will be conducted on several animal model in the future regarding graphene activity on wound closure. *In vitro* assays on endothelial cells may also help to understand if angiogenesis is facilitated in presence of graphene, or if the beneficial effects reported here are only due to an early downregulation of the inflammation phase.

## 5. Conclusion

Photographic wound assessment paired with histological analysis of skin and subcutaneous tissue show an enhanced healing in acute and chronic models with the use of a single layer of graphene dressing in direct contact with the wound bed. We introduced for the first time the use of CWT variation as a non-invasive, objective, and quantitative measurement of wound healing. Application of graphene dressing lessens the variation of temperature at the wound site during dressing removing. As the skin temperature is closely related to the blood flow underneath and inflammation process, our data strongly suggest that graphene improves early vascularization of the wound or shorten the inflammation phase. This hypothesis is furthermore supported by the results of the histological assessment in the acute model. This reduction of the inflammation phase may result from a graphene conductive scaffold property. This study also emphasized that the visual quantification by measuring wound area is not the most objective way to properly evaluate wound healing and other technics such as thermography should be implemented.

## ASSOCIATED CONTENT

SEM image of a fibroblast’s filipodium on graphene without metallization. Immunofluorescence of cells stained for DAPI, vimentin and actin in the absence or in the presence of graphene. Cytotoxicity study on graphene dressings in acute wound model. Schematic representation of the dressings used.

## AUTHOR INFORMATION

### Author Contributions

The manuscript was written through contributions of all authors. All authors have given approval to the final version of the manuscript.

### Notes

Vincent Bouchiat, Antoine Bourrier and Charlotte Hurot are involved in Grapheal SAS, a privately funded company focusing on biomedical applications of graphene.

## ACKNOWLEDGMENT

The authors thank Romaric Larcher, Yann Dumont and Sylvain Godreuil for their help screening the mouse skin flora.

## ABBREVIATIONS

CWT: cutaneous woud temperature
CVD: chemical vapor deposition
SEM: scanning electron microscopy
HMDS: hexylmethyldisilazane
BSA: bovine serum albumin
PFA: perfluoroalcoxy
DAPI: diamidino-2-phénylindole
TRITC: tetramethylrhodamine
PBS: phosphate buffer saline
DMEM: Dulbecco’s modified eagle medium
AFA: Alcohol-Formol-Acetic acid
CRP: C-reactive protein
CK: creatine kinase

## Notes

https://nextcloud.grenoble.cnrs.fr/s/ZZwcYT9kiXrzy59

### Competing Interest Statement

Vincent Bouchiat and Charlotte Hurot are involved in Grapheal, a privately funded company focusing on biomedical applications of graphene.

